# Social immunity as a driver of life-history evolution in eusocial species

**DOI:** 10.64898/2026.04.09.717360

**Authors:** Samir I. Aisin, Regina A. Belova, Denis A. Dmitriev, Peter V. Lidsky

## Abstract

Eusociality is accompanied by puzzling lifespan phenotypes that challenge classic theories of aging. In eusocial species, breeders age more slowly than non-breeders while sharing the same genomes. A notable exception is the naked mole-rat, in which all castes show negligible actuarial senescence. We show that both patterns can be explained with a single epidemiological model. Chronic parasites that reduce worker productivity can drive the evolution of shorter lifespan in workers, but not in queens. A genetic program that triggers the death of infected workers can evolve as an efficient alternative strategy for controlling pathogens, thereby reducing selection for shorter lifespan. However, in the presence of benign pathogens, this program results in excessive deaths and becomes too costly. Therefore, the composition of the pathogen mixture defines optimal life histories in eusocial communities: species exposed to a broad pathogen repertoire evolve caste differences in lifespan, whereas species occupying pathogen-poor environments are predicted to die rapidly upon infection and experience negligible aging. This framework links social immunity to life-history evolution and yields testable predictions for the pathogen control hypothesis of aging.

---

Eusociality evolved several times independently in colonial insects, crustaceans, and rodents. Eusocial societies comprise reproductive breeders and non-reproductive workers. Lifespan evolution in eusocial animals is a long-standing paradox: breeders typically age much more slowly and live much longer than worker castes, despite having identical genomes^1^. Additionally, the only mammal with strong evidence for negligible senescence, the naked mole-rat (*Heterocephalus glaber*), is an eusocial species ^2^. Both the longevity skew between breeders and workers and the negligible aging in naked mole-rats pose a challenge to simple versions of classic evolutionary theories of aging^3,4^.

Classic theories explain these phenotypes through trade-offs between early-life fitness and late-life survival, shaping life histories under different extrinsic hazard rates ^5,6^. However, these models remain incomplete because the nature of the trade-offs and the reasons for their evolutionary stability remain obscure^7^. Moreover, they do not fully account for the extreme lifespan plasticity of eusocial insects: workers can substantially extend their lifespans if they become reproductively active^8^, when manipulated by parasites^9^, or in response to colony size and social context^10^. It remains unclear why the longevity trajectory is not used ubiquitously, and a shorter lifespan and death from aging are selected instead^11,12^. Also, the negligible aging in naked mole-rat workers, whose annual mortality was estimated at 39%^13^, remains unexplained by the classic models.

## Results and Discussion

Here, we present an alternative framework of life-history evolution in eusocial communities. Unlike standard evolutionary accounts that treat aging as a byproduct of declining selection, we consider limited worker lifespan an adaptive trait that evolved to control the spread of chronic infectious diseases^14,15^ in the context of social immunity^16-18^. Evolutionary simulations showed that either a skew in lifespans between queens and workers, or negligible aging, can be shaped by the properties of pathogens and types of hosts’ immune reactions.

The life-history structure of our model communities was inspired by honeybee (*Apis mellifera*) colonies^19^ with some simplifications (Fig. 1a; Table S1). One colony consisted of a single reproductive queen and two thousand workers. Workers were further subdivided into nurses and foragers. Newborn workers assumed nursing roles, feeding larvae and cleaning the hive, and, upon this phase, transitioned to foraging. Foragers, who leave the hive to collect food, were subject to death due to extrinsic mortality (0.134 per day ^12^) or to an intrinsic lifespan limit, modeled as death upon reaching a specified age.

**Fig. 1.**
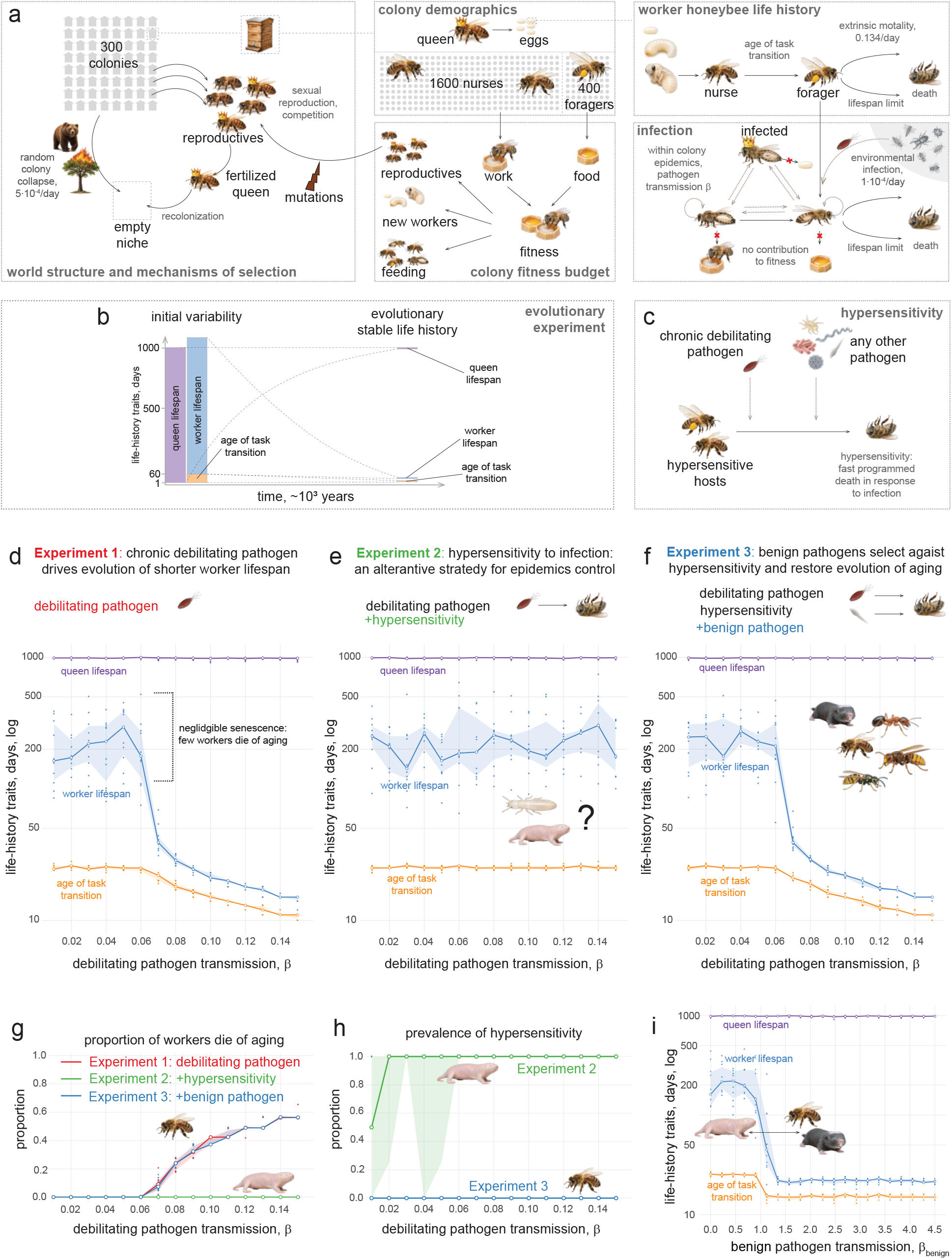
Modeling the life-history evolution in eusocial communities in the context of epidemics. **a**. The model’s structure. Colonies are subject to random collapse, and the emptied niches are filled by the reproductives coming from the remaining colonies. Each colony consists of a queen and workers: nurses and foragers. Workers contribute to the colony’s fitness budget, which is used to create new workers and feed the existing ones. The remaining fitness budget is invested in reproductives that determine the colony’s reproductive success. The newborn workers begin as nurses, and at the age of task transition, become foragers. Foragers are subject to dying due to extrinsic mortality or reaching an intrinsic lifespan limit (aging). In our model, infections that drive life-history evolution are chronic parasites that prevent workers from contributing to the fitness budget but do not increase their mortality. Such parasites first infect foragers and then spread within the colony. **b**. The evolutionary experiment. The initial allele variability was set in ranges [1:1000] for the queen’s lifespan; [1:60] for the age of the worker’s task transition; [1:1000] for the residual foragers’ lifespan. Workers’ cumulative lifespan was calculated as a sum of the last two values. The genotypes were allowed to compete for 1000 years. **c**. Hypersensitivity as a strategy to control epidemics: detection of infection triggers the host’s death. **d**. Chronic debilitating pathogens select for shorter workers’ lifespan and earlier task transition, but do not affect queens’ lifespan, resulting in the lifespan skew between castes. Medians of 10 independent replicates are shown here and in panels **e-i**. The colored areas are delimited by the 25th and 75th percentiles. **e**. Hypersensitivity can help control pathogen spread, making a limited lifespan redundant. **f**. High abundance of benign pathogens that trigger hypersensitivity reactions but do not incur substantial costs on their own selects against hypersensitivity and restores selection for a finite worker lifespan. **g**. The proportion of workers dying of aging with the evolutionary optimal life-histories found in **d-f. h**. The prevalence of hypersensitivity in the experiments described in **e** and **f. i**. The prevalence of a benign pathogen determines the transition between the two patterns of life histories in eusocial animals: skewed lifespan is predicted in pathogen-abundant environments, and hypersensitivity to infection combined with negligible senescence, in relatively isolated ones.

The world contained 300 colonies. The colony could collapse either after the death of all colony members or because of stochastic extinction events occurring with a probability of 5×10^-4^ per day. The emptied niches were filled by a new colony, whose genetic makeup was defined by the remaining colonies’ genotypes and their investment in reproductives: virgin queens and drones (Fig. 1a).

Colony output (fitness) was allocated to producing new workers, maintaining existing workers, and producing reproductives that compete to repopulate the freed niches. For tractability, colony fitness was calculated as a function of the healthy nurses and foragers, capped at 1600 and 400, respectively. A lower number of workers resulted in lower colony fitness, whereas an increased number did not yield further benefits but incurred additional costs for maintaining extra workers.

Three life-history traits were allowed to evolve: (i) age of task transition from nurse to forager, (ii) the overall worker lifespan, and (iii) the queen’s lifespan. Initial allele variability and *de novo* mutations were modeled (Fig. 1a, b; see Methods for details). We assumed sexual reproduction with codominant inheritance of the traits; phenotypes were computed as averages of the allele values.

Initially, we modeled epidemics of chronic parasites that do not kill workers but instead manipulate their behavior so that they stop working and contributing to the colony’s fitness as described in some species of eusocial insects^9,20-22^. Infected queens were considered sterile^23^. We modeled the scenario in which each forager may contract the parasite from the environmental source^21^ with a constant probability of 10^-4^ per day and later infect other colony members with a variable probability of β (pathogen transmission efficiency). The value of β determined pathogen prevalence and thus, the selection pressure exerted by pathogens on their hosts. No recovery from infection was assumed.

First, we conducted a series of evolutionary experiments in the presence of pathogens with varying β values (Fig. 1d). At low transmission, 0.0 ≤ β ≤ 0.04, pathogens were not able to propagate, and the worker’s lifespan fixed above 100 days, which corresponded to effectively negligible mortality from intrinsic reasons, as it caused death in a minimal proportion of workers (Fig. 1g). The age of task transition evolved towards ∼26 days to ensure an optimal demographic balance in the colony.

However, with β ≥ 0.05, a younger age of task transition and a shorter worker’s lifespan became evolutionarily beneficial. The rapid mortality of workers helped remove potentially infected individuals who might otherwise have spread the disease within the colony and diminish its fitness. This prevention of epidemics (Fig. S1) came at the cost of a higher proportion of animals dying from aging (Fig. 1g). Contrastingly, the evolution of queens’ lifespan was unaffected by pathogens (Fig. 1d), because rapid worker turnover protected queens from infection. Therefore, according to our model, the skew in the life histories of eusocial queens and workers can be determined by selection for social immunity.

However, mortality at older age is not the only strategy for controlling epidemics^24^. Under certain conditions, programmed self-killing in response to infection may evolve to block onward transmission. Such programmed death mechanisms in response to infection are present in bacteria^25^. In eusocial workers, altruistic responses to infection are well established and are widely discussed in the context of social immunity^26,27^. The mechanisms include self-isolation^27,28^, self-destruction^29^, or destruction by other colony members^30-32^ .

To test whether a self-destruction in response to infection can be evolutionarily stable in eusocial communities, we introduced a dominant “hypersensitivity” allele that increased workers’ daily mortality up to 0.5 if they were infected (Fig. 1c). We found that under pathogen pressure, the hypersensitivity allele was fixed in eusocial populations, resulting in efficient control of infection. Moreover, in the presence of hypersensitivity, the short lifespan limit became redundant as a mechanism for epidemic prevention (Fig. 1e), resulting in negligible numbers of individuals dying of intrinsic reasons, independent of the presence of chronic pathogens (Fig. 1g). Queens’ lifespan was fixed at its maximum possible value as in the previous case (Fig. 1e).

If pathogens are indeed a selective force shaping life-history evolution in eusocial communities, hypersensitivity to infection can become a viable and preferable alternative to a shorter lifespan. This can explain the negligible actuarial senescence in naked mole-rats^2^. Indeed, some indirect evidence suggests that the naked mole-rats may be hypersensitive to infections. In particular, they die shortly after inoculation with herpes simplex virus^33^ (which is sublethal in mice and used for gene delivery^34^), are susceptible to lethal viral epidemics^35^, and have a unique immune system architecture^36^ consistent with the unusual reaction to infections. Of note, the naked mole-rat workers show faster aging, according to the methylation clock readouts^37^, compared to queens, in good agreement with our model. Testing whether naked mole rats possess immune mechanisms of self-killing in response to infection would provide a means of evaluating the predictions of our model.

However, if a combination of hypersensitivity and negligible senescence is a better strategy to control epidemics in eusocial communities, why did it not evolve ubiquitously? According to our model, the reason is the composition of the pathogen mixture. If the community is exposed to multiple infections, not all of which are highly debilitating, the cost of hypersensitivity as a protective mechanism may become prohibitive. The adaptive killing program will be triggered by benign infections that impose minimal fitness costs. Thus, high prevalence of such benign pathogens can drive selection against hypersensitivity.

To test this hypothesis, in addition to a chronic debilitating pathogen, we introduced a second, benign pathogen with a very high transmission rate (β_benign_ = 4.5), without any direct cost to animal fitness, but capable of triggering the host’s hypersensitivity response (Fig. 1c). Under these conditions, hypersensitivity was rapidly lost and, as a result, selection toward limited worker lifespan and adaptive lifespan limit was restored, matching the results shown in Fig. 1d.

These results indicate that the composition of the pathogen mixture, specifically the abundance and fitness penalties that pathogens impose on their hosts, may determine the evolutionary balance between individual and social immunity and the life histories in eusocial communities. In pathogen-poor environments where the risk of infection is low, eusocial animals evolve hypersensitivity to control a few remaining chronic pathogens and negligible aging. In addition to the naked mole-rats, the same pattern can be predicted in eusocial insects whose nests are protected from infection. In such niches, the model predicts negligible senescence, low viral loads, rapid death after infection, and abnormal immune responses. Potential candidates can be deep underground or tree-dwelling termite species^38^. On the contrary, eusocial communities exposed to environments abundant in pathogen species are expected to evolve tolerance to infections and life-history patterns with short-lived workers and long-lived queens, as seen in most eusocial insects^1^ and Fukomys mole-rats^39^. Importantly, when comparing the lifespans of different worker castes, the infection hazard is predicted to play a larger role than extrinsic mortality risk^5^. Additionally, a more complex combination of individual and social immune mechanisms can evolve in pathogen-abundant environments. For example, ants of the species *Lasius niger* rely on social immune responses to fungal infections and on individual immunity to bacterial diseases^40^.

In sum, our model identified the key epidemiological parameters, the composition of the pathogen mixture, transmission, and cost of infection, that explain puzzling patterns associated with aging in eusocial species. Critically, we demonstrate that changes in pathogen abundance can generate transitions between queen- and worker-skewed and all-caste negligible aging (Fig. 1i).

Therefore, the life-history evolution of eusocial animals can be explained by combining the theoretical framework of social immunity^18,24^ with the pathogen control theory of aging^7,14,15^. Importantly, our explanation neither contradicts nor requires the classic antagonistic pleiotropy arguments^5^. Further experimental validation can test our predictions concerning hypersensitivity, pathogen loads across different eusocial species, and the correlation between caste-specific longevity and infection hazard.

## Model description

### World structure

We modeled 300 colonies with 2001 productive members: 1 queen and 2000 workers.

### Initialization

For each colony, a random genotype was generated. The diploid genome contained 4 alleles: queen lifespan (in an interval 1:1000), age of task transition (1:60), and forager lifespan after task transition (1:1000). In experiments 2 and 3, the hypersensitivity alleles were present in half of the queens. Initially, all queens were generated as homozygotes. At the beginning of the simulation, each animal is assigned a random initial age within a range allowed by its genome.

### Colony’s fitness

Each day, a colony’s fitness is calculated as a product of the number of uninfected nurses and foragers, capped at 1600 and 400, respectively. Next, the costs of producing new workers (10^3^ fitness units per worker) and feeding the existing colony members (30 fitness units per animal per day) were subtracted. The residual amount of fitness was invested into the reproductives and used as metrics of the colony’s reproductive success.

### Newborns

If a queen is alive and fertile, it produces eggs, which develop into newborn workers. For simplicity, we assumed instant development as a space in the colony becomes vacant, with the maximum number of newborns capped at 200 per day.

### Task transition

After a nurse reaches its task transition age, it becomes a forager.

### Extrinsic and intrinsic death

The extrinsic mortality concerned only foragers and was assumed to be 0.134 per day. For ease of calculation, the intrinsic mortality in foragers and queens was modeled as an immediate death upon reaching the critical age.

### Colony collapse

Each colony has a daily probability of collapse due to extrinsic reasons of 5×10^-4^. Additionally, colonies were considered extinct if all their members were dead and the queen was sterile. The queen’s death led to the colony’s collapse; no queen replacement was modeled. Colony death emptied a niche for recolonization.

### Replication

Each empty niche was recolonized by a queen fertilized by a drone. Both parents originated from the remaining colonies. The probability of participating in recolonization was proportional to the colony’s reproductive investment. Therefore, for each gene, one of the two alleles is inherited from each parent.

### Genetics

The inheritance of life-history traits was codominant: the phenotype was determined by averaging the effects of the two alleles, with the hypersensitivity allele considered dominant. For simplicity, we assumed the queen’s lifespan to be determined by their progeny’s genomes.

### Mutations

The mutations occurred with the probability of 0.1 per recolonization event. One mutation affected only one allele and changed its value from 1 to 1000. Mutations that shorten lifespan or accelerate task transitions occurred more often than those that extend it. New values upon mutation were calculated according to the formula: *N* = 10^(*logA*+*X*·(*logB*−*logA*))^, where N is the new value of the allele; A and B are the lowest and the highest possible values, respectively; 0≤X≤1 – a random value.

### Infection

There were two types of pathogen transmission: infection spilled from the environment and the within-colony infection. The environmental spill affected only foragers with a constant probability of 1×10^-4^ per day. The within-colony transmission occurred with equal probability among all colony members. The probability of infection of each susceptible colony member was calculated as *I* · *β*, where I is the number of infected colony members, and *β* is the pathogen transmission efficiency. No infection was assumed between different colonies, and no recovery from the infection was modeled.

### Two types of pathogens

We modeled two types of pathogens: a debilitating pathogen that abolishes the worker’s contribution to fitness, and a benign pathogen that does not affect workers by itself but can activate their hypersensitivity responses. debilitating pathogen. The benign transmission (*β*) was studied in the interval from 0 to 0.15 (Fig. 1d-f) and was set to 0.1 in panel 1i; while the benign pathogen transmission (*β*_*benign*_) in the interval from 0 to 4.5 and set to 4.5 in panel 1f.

### Hypersensitivity

The hypersensitivity allele increased the risk of death (up to 0.5/day) in nurses and foragers.

### Termination of the simulation

The simulations were run until either the evolutionary steady state was reached or the time limit was reached. The criteria for the steady state were low variability among the three life-history allele values (queen lifespan, age of task transition, forager lifespan) across all 300 colonies (coefficient variation <0.1) and the evolutionary stability of these values over 900 days (change <0.1). The time limit was set to 10^3^ years (3.65·10^6^). If one of these conditions holds, the simulations were terminated and the resulting values recorded.

## Data collection and presentation

At the end of each simulation, the values of the surviving alleles for life-history traits, the proportion of dead workers in the last day, and the proportion of hypersensitivity alleles were collected from all 300 hives to produce medians. 10 replicates per experiment were used. The values presented in Fig. 1 are medians and percentiles derived from these replicates.

## Coding

The random probabilities were generated with the xoshiro256++ algorithm. The simulation is written using programming languages: Mojo (version 0.26.1), Python (version 3.11), R (version 4.4.3, http://www.r-project.org).

## Data availability

The code can be downloaded from https://github.com/DivaythFyr/collective_immunity_shapes_aging_in_eusocial_Supplementary/

## Model limitations

The model does not account for behavioral, demographic, and life-history plasticity. Eusocial animals can change their behaviors, life-history, and social network structures^41^ in response to environmental factors, including infections. However, we believe that the interactions identified by our models are fundamental, and the conclusions will stand in the presence of plasticity in response to infection.

---

**Fig S1.**
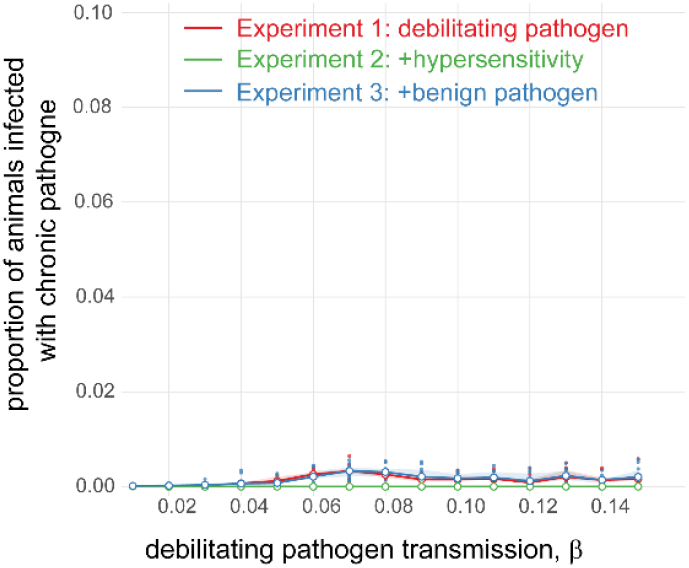
Low prevalence of the debilitating pathogen across three experiments (Fig 1d-f). Under the assumptions used, the evolutionary stable life histories led to an efficient prevention of epidemics of debilitating disease.

**Supplementary Table 1.** Model parameters.

| Variables | Meaning | Values | Unit | Comments |
| --- | --- | --- | --- | --- |
| $m$ | Mutation probability | 0.1 | per replication | |
| $\mu$ | Forager extrinsic mortality | 0.134 | day<sup>-1</sup> | mortality rate is from<sup>12</sup> |
| $D$ | Probability of colony death | 0.0005 | day<sup>-1</sup> | Colony's extrinsic destruction |
| $T_{max}$ | Maximum simulation time | $3.56 \cdot 10^5$ | day | |
| $A$ | Maximum number of animals per colony | 2001 | units | Reduced to optimize performance |
| $N$ | Optimal number of nurses | 1600 | units/colony | 80:20 estimate is from<sup>19</sup> |
| $F$ | Optimal number of foragers | 400 | units/colony | |
| $Q$ | Number of queens | 1 | units/colony | |
| $E$ | Maximum number of eggs hatching | 200 | units/day | |
| $W$ | Colony fitness | variable | fitness units | N·F |
| $C$ | Fitness cost of producing a worker | 1000 | fitness units | |
| $p$ | Maintenance cost for a colony member | 30 | fitness units/day | |
| $B$ | Probability of environmental infection | 0.0001 | units<sup>-1</sup>day<sup>-1</sup> | Constant |
| $\beta$ | Transmission of a debilitating pathogen | [0.0,0.15] | units<sup>-1</sup>day<sup>-1</sup> | varied between simulations |
| $\beta_{benign}$ | Transmission of a benign pathogen | [0.0,4.5] | units<sup>-1</sup>day<sup>-1</sup> | varied between simulations |
| $h$ | Probability of death due to hypersensitivity | 0.5 | day<sup>-1</sup> | in infected individuals only |

